# Simultaneous mnemonic and predictive representations in the auditory cortex

**DOI:** 10.1101/2021.10.08.463638

**Authors:** Drew Cappotto, HiJee Kang, Kongyan Li, Lucia Melloni, Jan Schnupp, Ryszard Auksztulewicz

## Abstract

Recent studies have shown that stimulus history can be decoded via the use of broadband sensory impulses to reactivate mnemonic representations. It has also been shown that predictive mechanisms in the auditory system demonstrate similar tonotopic organization of neural activity as that elicited by the perceived stimuli. However, it remains unclear if the mnemonic and predictive information can be decoded from cortical activity simultaneously and from overlapping neural populations. Here, we recorded neural activity using electrocorticography (ECoG) in the auditory cortex of anesthetized rats while exposed to repeated stimulus sequences, where events within the sequence were occasionally replaced with a broadband noise burst or omitted entirely. We show that both stimulus history and predicted stimuli can be decoded from neural responses to broadband impulse at overlapping latencies but linked to largely independent neural populations. We also demonstrate that predictive representations are learned over the course of stimulation at two distinct time scales, reflected in two dissociable time windows of neural activity. These results establish a valuable tool for investigating the neural mechanisms of passive sequence learning, memory encoding, and prediction mechanisms within a single paradigm, and provide novel evidence for learning predictive representations even under anaesthesia.

## Introduction

The presence of multiple stimulus streams in natural environments presents a unique challenge to our perceptual systems, as monitoring all of these streams in real time to extract useful information would require a tremendous amount of cognitive resources that our brains might otherwise use in service of other tasks. The predictive coding framework proposes one possible way in which our brains deal with this challenge (Friston, 2005; Heilbron and Chait, 2018). Stimulus streams tend to have consistent, repetitive features. Sensory systems thus do not necessarily need to constantly parse an endless stream of incoming information, but can instead make this process more efficient by forming predictions based on the regularities of previously observed streams. These predictions allow the brain to create a probable model of the outside world, which can be updated when errors are detected between the model predictions and external inputs (Fairhall et al., 2001; Friston et al., 2006; Rubin et al., 2016; Schröger et al., 2014).

In the auditory system, explanations based on predictive coding have been applied to several phenomena, such as temporal expectation, mismatch negativity (MMN) responses, and so-called omission responses (Denham and Winkler, 2020; Heilbron and Chait, 2018). In all of these contexts, experimental paradigms typically employ the use of repetitive stimulus streams which allow a build-up of predictions, based on memory of recent stimulation. Within the predictive coding framework, memory is intrinsically linked with predictive mechanisms in the form of adaptive memory traces employed in downstream error correction (Wacongne et al., 2012). However, memory and prediction formation can also be studied more directly under sequence learning paradigms. Research has demonstrated the brain’s ability to recognize sequential patterns independently of attentional state (Denham and Winkler, 2020), with previous studies establishing neural markers of sequence learning across species (Henin et al., 2021; Kikuchi et al., 2018, 2017), suggesting the underlying mechanisms are also evolutionarily conserved.

In the context of sequence processing, “mnemonic representations” would entail a memory of past sequence elements, independent of the currently processed sequence element, while “predictive representations” would entail a specific prediction of which sequence element is expected at a given time, as well as the comparison of this prediction with the currently processed stimulus. These representations are intrinsically linked, as prior mnemonic representations service the accrual of information across previously learned patterns, while predictive representations can be seen as a form of memory retrieval used to predict the current or future sensory states. Indeed, sequence learning has been shown to mediate predictive mechanisms in sensory cortices (Luft et al., 2015) in addition to prefrontal areas (Henin et al., 2021; Kikuchi et al., 2017). However, auditory cortical activity has also been shown to mediate working memory maintenance, with deactivation of cortical neurons during stimulus presentation significantly impairing memory task performance (Yu et al., 2021). Recent work has suggested possible cortical (and cortico-hippocampal) mechanisms for encoding mnemonic and predictive representations in the same regions, but using distinct neuronal populations (Barron et al., 2020). Furthermore, direct recordings in the auditory cortex of awake mice established neural mechanisms for how encoding mechanisms might handle working memory and predictive processes without “overwriting” recent sensory events in instances where predictive mechanisms are triggered by oddballs within a sequence (Libby and Buschman, 2021). Nevertheless, the implementation of paired memory and predictive mechanisms remains a largely unexplored topic ripe for many lines of investigation. Memory and prediction have traditionally been tied to attention, in that both phenomena typically are studied in contexts where information is task-relevant (Libby and Buschman, 2021, Doron, et al, 2020). However, it has recently been argued that working memory contents can be encoded both as active states (selected to guide imminent behaviour) and as passive states (stored for potential later use; Muhle-Karbe et al., 2021). Similarly, both active states (such as top-down suppression) and passive states (such as short-term plasticity) have been implied in predictive processing (Garrido et al., 2009; Auksztulewicz et al., 2017; van Moorselaar & Slagter, 2019). It remains to be understood whether both mnemonic and predictive representations can be encoded in parallel without any attentional involvement (e.g., under anaesthesia), and whether their passive encoding relies on dissociable neural codes, as recently observed for active states (Libby and Buschman, 2021).

The informational state of working memory networks can be accessed using broadband noise bursts (Cappotto et al., 2021; Stokes, 2015; Wolff et al., 2020, 2015), which are thought to reactivate the activity-silent representation stored in synaptic weights. Recent work in the decoding of auditory sensory memory traces across species offers support for the notion of evolutionarily conserved memory encoding mechanisms (Cappotto et al., 2021). Conversely, predictive mechanisms are most commonly investigated through the use of occasional, and unpredictable, sensory deviants or omissions of stimulus tokens within a repeated sequence. The neural responses to these events contain information about the identity of the predicted token that is unconfounded with responses to perceived stimuli. This phenomenon has been demonstrated for predicted pitch contours as well as for individual token values omitted from repeated sequences (Berlot et al., 2018; Chouiter et al., 2015; Demarchi et al., 2019). However, sequence learning paradigms combined with occasional presentations of broadband noise bursts (a causal means to probe the state of the network) offer a possibility to tap into the mnemonic and predictive representations using the same methods in a single paradigm.

Here we adapt recent techniques for the decoding of auditory working memory traces to simultaneously probe both phenomena. Electrocorticography (ECoG) was recorded from the auditory cortex (AC) of anesthetized rats (N=8) while repeated stimulus streams of artificial vowel tokens were played (referred to henceforth as “vowels”, for simplicity). The streams were occasionally interrupted where vowels were replaced by either a burst of broadband noise or omitted entirely, and multivariate analysis was employed to investigate the decodability of stimulus history as well as predictive mechanisms. This approach allows us to leverage an easily-implemented decoding paradigm to investigate what have traditionally been discrete research questions, establishing a new model for simultaneous memory and prediction research.

## Results

### Auditory cortical activity recordings in a sequence learning paradigm

To investigate the decodability of mnemonic and predictive representations from AC activity, we combined ECoG recordings in young adult female Wistar rats (N = 8) with an auditory sequence learning paradigm. All rats were “naive”, i.e. had no experience with the stimulus sequences prior to recording, and anaesthetised before being implanted with ECoG electrode arrays over their auditory cortex and exposed to stimulus sequences (Fig. 1A; see Materials and Methods for details on experimental procedures). Frequency Response Area (FRA) maps for each animal were used to visually verify whether the placement of the array was consistent across subjects (Fig. 1B).

**Figure 1.**
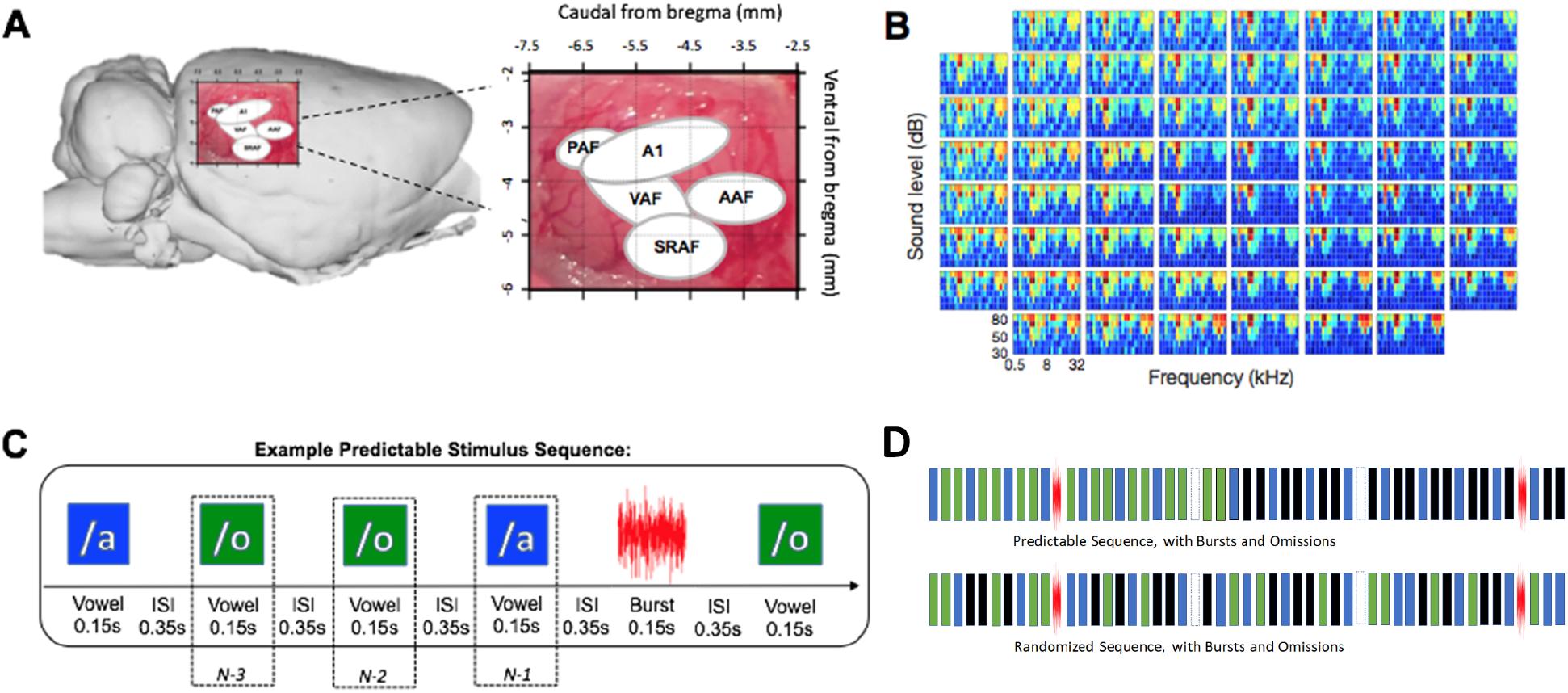
Electrocorticography methods and acoustic stimulation. **(A)** Electrode placement during electrocorticography. **(B)** An example Frequency Response Area map from one subject. These were used to confirm electrode array placement consistency over the AC across subjects. **(C)** An example of an AOO predictable stimulus sequence, where one vowel of the triplet has been randomly substituted by a noise burst (or, not shown, alternately omitted entirely) following a minimum of three triplet repetitions. In paired random blocks, the relative position of the burst/omission substitution remains unchanged, while the surrounding vowel vowels are randomized. Vowel positions relative to the burst/omission are denoted as N-1, N-2, and N-3. **(D)** Segment of an example predictable sequence, in which vowel tokens are omitted or replaced with a noise burst after 3 repetitions (top) and the randomized version of that sequence where vowel tokens from the full sequence are presented pseudo-randomly while burst and omission tokens remained in the same relative positions (bottom).

The sound stimuli (Fig. 1C) consisted of sequences of artificial vowels of 150 ms in duration, separated by 350 ms of silence (500 ms onset to onset ISI). Three artificial vowels (A, I, O) were used in the experiment. We deemed artificial vowels preferable to click trains as they activate larger parts of the tonotopic array and are arguably more ecologically valid than pure tones. Sequences were presented in two experimental conditions, across separate blocks. In “predictable” blocks, vowels were grouped into triplets, which repeated at least 25 times (range 25-100, mean 30) before being replaced with another triplet. In “random” blocks, vowels were presented in a random order, while keeping the base frequency of each vowel constant and comparable to the predictable block. In both types of blocks, 5% of vowels were replaced with omissions, and 5% were replaced with a burst of broadband noise.

### Univariate analyses: only vowel-evoked activity differentiates between vowels

First, to test whether vowel identity influences mean neural activity in the AC at a coarse spatial resolution (i.e., on a channel-by-channel basis), we have performed a series of univariate analyses, testing for the effects of vowel (A, I, O) and block (predictable vs. random) on vowel-evoked ECoG activity (event-related potentials). We observed that vowel-evoked activity does differentiate between the three vowels, both in predictable blocks (Fig. 2A; 13-260 ms; F_max_= 58.56; p_FWE_ < 0.001) and in random blocks (Fig. 2B; 13-207 ms; F_max_ = 58.21; p_FWE_ < 0.001). The main effect of block on vowel-evoked activity was not significant (all p_FWE_ > 0.05).

**Figure 2.**
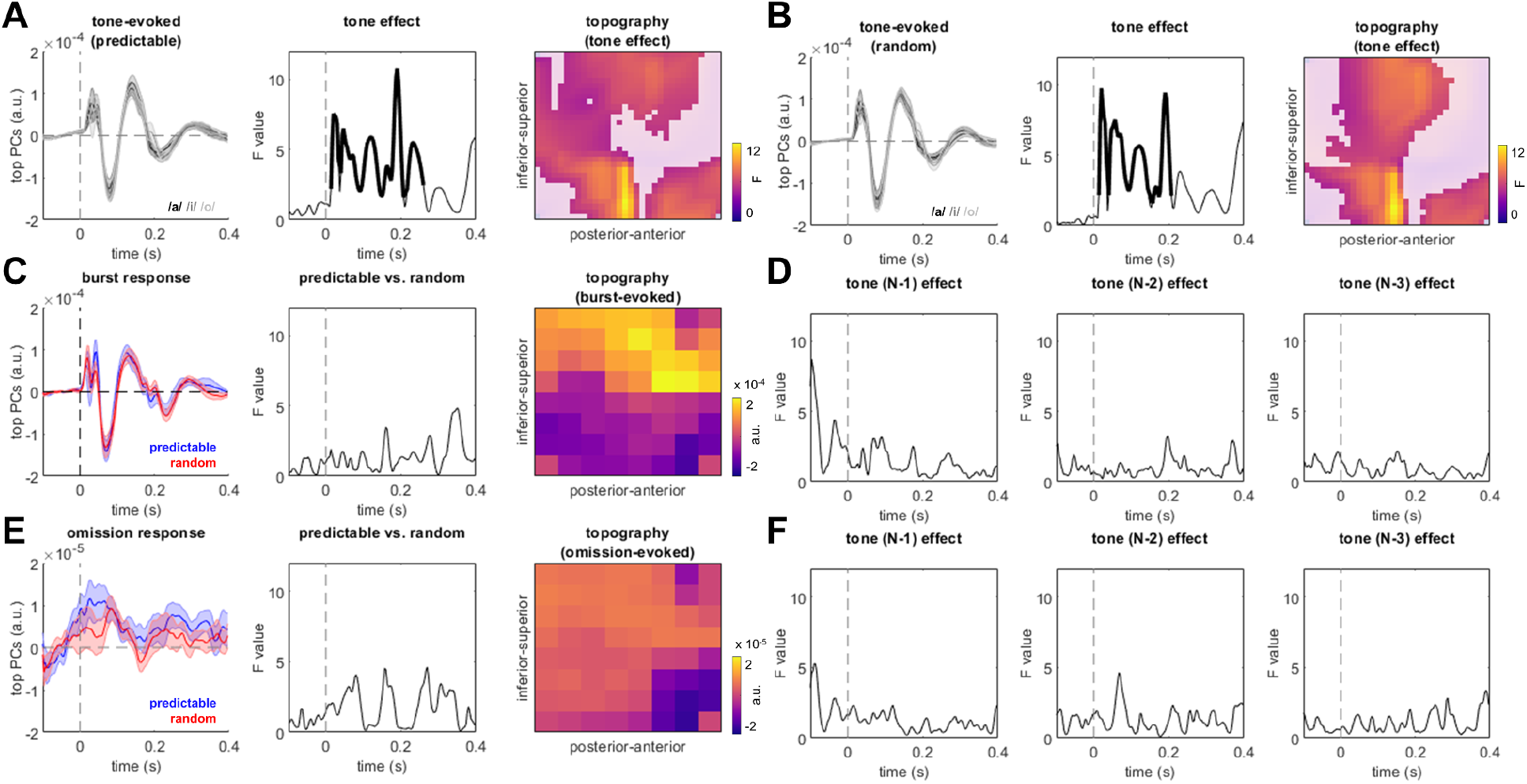
Univariate analyses. **(A)** vowel-evoked responses in predictable blocks. Left panel: Time courses of vowel-evoked responses, summarising the top principal components explaining 95% of the variance (shaded area: SEM across recording sessions). Middle panel: Time course of the main effect of vowel (bold: p_FWE_ < 0.05). Right panel: Topography of the main effect of vowel (unmasked area: p_FWE_ < 0.05). **(B)** vowel-evoked responses in random blocks. Figure legend as in (A). **(C)** Burst-evoked responses. Left panel: Time courses of noise burst-evoked responses, summarising the top principal components explaining 95% of the variance (blue: predictable blocks; red: random blocks; shaded area: SEM across recording sessions). Middle panel: Time course of the main effect of predictability. No significant differences were observed between predictable and random blocks. Right panel: Topography of the noise burst-evoked responses, averaged across blocks and recording sessions, summarising the top principal components explaining 95% of the variance. **(D)** Effects of preceding vowel on noise burst-evoked responses. Left/middle/right panel: Time courses of the main effect of the preceding vowels (N-1 / N-2 / N-3, relative to noise burst). No significant differences were observed. **(E)** Omission-evoked responses. Figure legend as in (C). No significant differences were observed between predictable and random blocks. **(F)** Effects of preceding vowel on omission-evoked responses. Figure legend as in (D). No significant differences were observed.

Having established that vowel-evoked activity differentiates between the three vowels, but not between experimental conditions (predictable vs. random), we then tested whether burst-evoked and/or omission-evoked activity also differentiates between the (preceding) vowels at different “positions” in the sequence, relative to the burst/omission (N-1 position: the immediately preceding vowel, N-2 position: two stimuli before the burst/omission, N-3 position: three stimuli before the burst/omission). This analysis revealed that, similarly to the vowel-evoked responses, burst-evoked responses did not significantly differentiate between predictable and random blocks (Fig. 2C; all p_FWE_ > 0.05). However, unlike the vowel-evoked activity (which was modulated by vowel identity), noise burst-evoked activity was not significantly modulated by (preceding) vowel identity when neural activity was analyzed in a mass-univariate manner. Specifically, neither the effect of the immediately preceding vowel on burst responses (N-1: all p_FWE_ > 0.05; Fig. 2D), nor of the previous vowels (N-2, N-3: all p_FWE_ > 0.05) were significant.

Omission-evoked responses peaked relatively early (83-93 ms), with a rising activity visible already prior to expected stimulus onset, and thus possibly marking the offset response to the preceding interrupted stimulus train rather than a true omission (Chien et al., 2019). Nevertheless, just like burst-evoked activity, omission-related activity was also not significantly modulated by block type (Fig. 2E; all p_FWE_ > 0.05) or preceding vowel identity (Fig. 2F; N-1, N-2, N-3: all p_FWE_ > 0.05).

### Univariate analyses: spectral peaks observed for single vowel rate but not triplet rate

In a frequency-domain analysis, we tested whether sequence processing is associated with frequency peaks in the neural response spectrum at the vowel (2 Hz) and triplet (0.66 Hz) rate, as reported in ECoG studies in humans (Henin et al., 2021). To this end, we analyzed phase coherence of neural activity and observed robust spectral peaks at the single vowel rate (Wilcoxon’s signed-rank test against neighbouring frequency points: Z = 5.6545, p < 0.001; Fig. S1A), but not at the triplet rate (p = 0.4569), consistent with a recent study in anaesthetised rats (Luo et al., 2021). No differences in spectral peaks were observed between predictable and random blocks at either the single vowel rate (p = 0.7943) or the triplet rate (p = 0.6639).

### Multivariate analysis: specific decoding boost for predictable vowels

Although noise burst-evoked activity did not differentiate between preceding vowels when analyzed in a mass-univariate way, based on our previous study (Cappotto et al., 2021) we hypothesized that preceding stimuli can nevertheless be decoded in a multivariate analysis.

Specifically, by analyzing the spatiotemporal pattern of activity evoked by noise bursts, which did not carry any overt information about the preceding vowels given that the employed noise tokens were always identical and presented after vowel-evoked responses had returned to baseline (400 ms after stimulus offset), we sought to determine if activity evoked by noise bursts contained information about the preceding vowels (separately for N-1, N-2, and N-3 vowels). This analysis revealed significant decoding of vowels up to N-3 (Fig. 3A) in predictable blocks (N-1: -10-273 ms, t_max_ = 13.36, cluster-level p_FWE_ < 0.001; N-2, early cluster: 40-106 ms, t_max_ = 3.83, cluster-level p_FWE_ < 0.001; N-2, late cluster: 166-203 ms, t_max_ = 3.23, cluster-level p_FWE_ = 0.001; N-3, early cluster: 33-123 ms, t_max_ = 5.44, cluster-level p_FWE_ < 0.001; N-3, late cluster: 223-240 ms, t_max_ = 2.98, cluster-level p_FWE_ = 0.044). Decoding was significantly better for the immediately preceding (N-1) vowels than for the earlier vowels (N-1 vs. N-2: -6-236 ms, t_max_ = 7.36, cluster-level p_FWE_ < 0.001; N-1 vs. N-3: -3-216 ms, t_max_ = 7.84, cluster-level p_FWE_ < 0.001). Clusters of significant decoding extended into the baseline are likely due to the sliding time window approach (50 ms) adopted in the multivariate analyses (see Methods).

**Figure 3.**
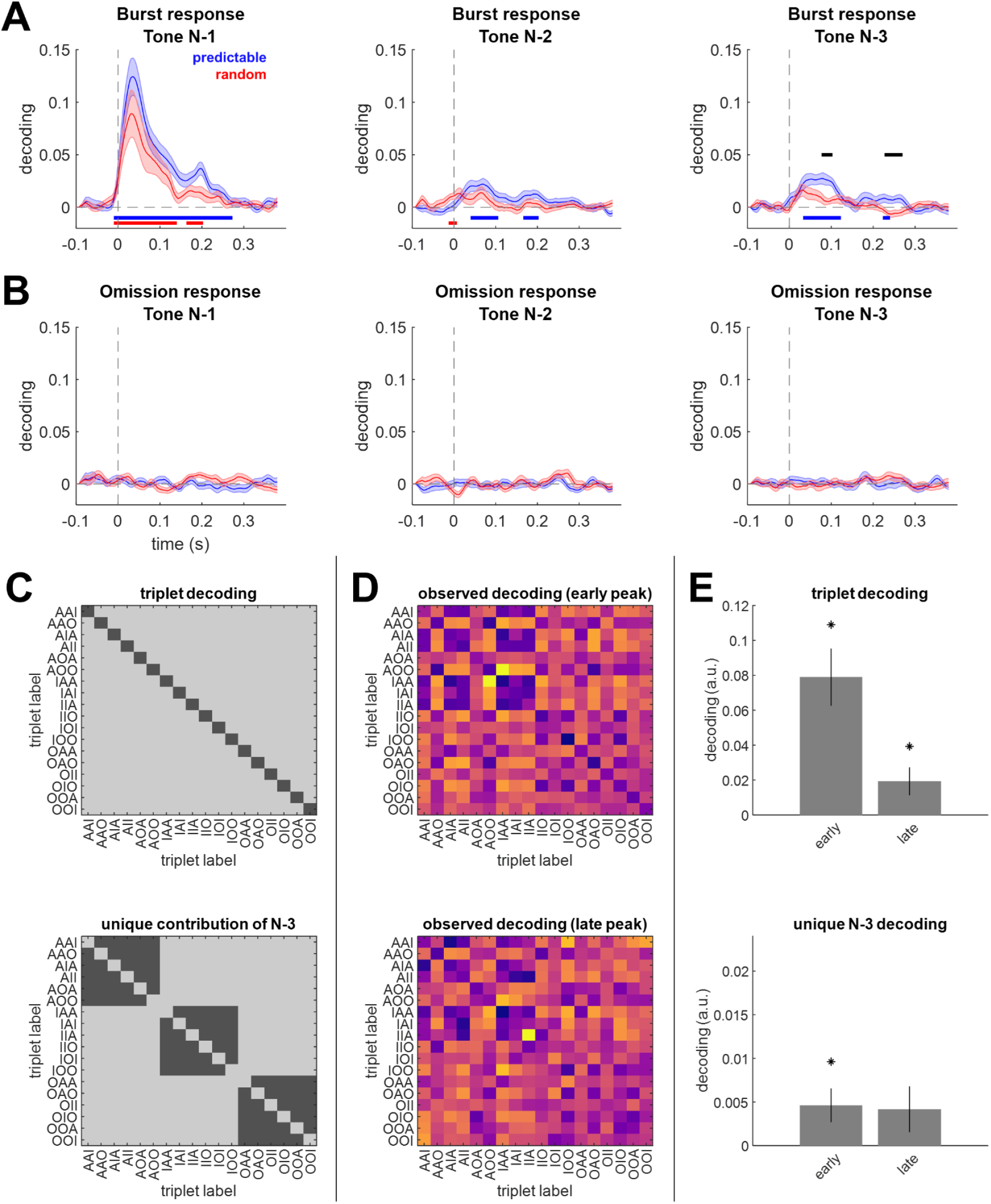
Multivariate analyses. **(A)** Time courses of decoding of preceding vowels based on burst-evoked activity. Left / middle / right panel: decoding N-1 / N-2 / N-3 vowel (blue: predictable blocks; red: random blocks; shaded area: SEM across recording sessions; blue/red horizontal line: decoding in predictable/random blocks significantly different from zero, p_FWE_ < 0.05; black horizontal line: decoding significantly different between predictable and random blocks, p_FWE_ < 0.05). **(B)** Time courses of decoding of preceding vowels based on omission-evoked activity. Figure legend as in (A). **(C)** Stimulus representational dissimilarity matrices used to quantify the decoding of entire triplets (upper panel) and the unique contribution of decoding the first element of each triplet (i.e., N-3 vowels) while excluding entire triplets along the diagonal (lower panel). Darker gray shows stimulus similarity, lighter gray shows stimulus dissimilarity. **(D)** Observed decoding matrices for the early peak (left panel) vs. late peak (right panel) observed in the N-3 decoding trace (A, right panel). “Warmer” colors denote larger Mahalanobis distance (a.u.). **(E)** Triplet decoding (left panel) and the unique contribution of decoding the first element of each triplet (N-3 vowel; right panel) for the early and late peaks (separate bars). Error bars denote SEM across recording sessions. Asterisks denote significance (p < 0.05).

In random blocks as well, significant decoding of preceding vowels was possible, but only the N-1 and N-2 vowels could be decoded (N-1, early cluster: -10-140 ms, t_max_ = 7.73, cluster-level p_FWE_ < 0.001; N-1, late cluster: 163-203 ms, t_max_ = 3.26, cluster-level p_FWE_ = 0.001; N-2: -13-7 ms, t_max_ = 3.03, cluster-level p_FWE_ = 0.029). As was the case for predictable blocks, in the random blocks decoding was also significantly better for the immediately preceding (N-1) vowels than for the earlier vowels (N-1 vs. N-2: 0-136 ms, t_max_ = 5.91, cluster-level p_FWE_ < 0.001; N-1 vs. N-3: -6-126 ms, t_max_ = 5.53, cluster-level p_FWE_ < 0.001). Thus, using multivariate analyses, we could access mnemonic representations reactivated by a noise burst, up to 3 stimuli in the past in the predictable blocks and up to 2 stimuli in the past in the random blocks. Overall, immediately preceding stimuli could be decoded better than previous stimuli.

Crucially, if burst-evoked activity can reactivate not only mnemonic representations (irrespective of the currently processed stimulus), but also predictive representations (regarding the stimulus which would have been predicted but is replaced by a noise burst), we would expect a specific decoding improvement for N-3 (but not N-2 or N-1) vowels presented in predictable blocks vs. random blocks. The decoding results were consistent with this hypothesis. Specifically, a significant decoding boost was only observed for the N-3 vowels presented in predictable blocks (early cluster: 77-103 ms, t_max_ = 3.45, cluster-level p_FWE_ = 0.010; late cluster: 227-270 ms, t_max_ = 3.79, cluster-level p_FWE_ < 0.001), suggesting that we could access a predictive representation of the vowel replaced by a noise burst.

While mnemonic and predictive representations could be decoded based on burst-evoked activity, decoding stimulus history based on omission-evoked activity did not yield any significant results (Fig. 3B; all p_FWE_ > 0.05). This suggests that, at least in this experimental protocol (vowel triplets) and under anesthesia, a stronger activation of the network (e.g., burst presentation) is necessary to make mnemonic and/or predictive representations observable in this type of extracellular recordings.

The finding that burst-evoked responses in predictable blocks could be used to decode mnemonic representations about the past 3 triplet elements, as well as predictive representations about the N-3 vowel, entails a possibility that what is being represented is the entire triplet, rather than unique information about the predicted N-3 vowel independently of the remaining triplet elements (N-2 and N-1). To test this hypothesis, we repeated the decoding analysis, this time decoding stimulus information on a triplet-by-triplet level (Fig. 3C). This analysis revealed that the entire triplet identity can indeed be decoded from burst-evoked responses, both for the early decoding cluster (77-103 ms; t_20_ = 4.8118; p < 0.001) and the later cluster (227-270 ms; t_20_ = 2.4075; p = 0.0258; Fig. 3DE). However, information unique to the N-3 vowel but independent of triplet identity was also present in the early decoding cluster (t_20_ = 2.3889, p = 0.0269), but not in the later cluster (t_20_ = 1.5881; p = 0.128; Fig. 3E). This suggests that a predictive representation of the expected triplet element is decodable shortly following the noise burst that replaces the expected vowel, while a representation of the entire triplet is present at a wider range of latencies following the noise burst.

In two supplementary analyses, we tested whether decodability primarily relies on those channels which are also associated with sensory encoding of vowels, and/or on those channels which show robust phase coherence at the syllable rate. The first analysis revealed a significant main effect of channel selection (17-40 ms; Fmax = 10.45; p_FWE_ < 0.001; Fig. S1B), suggesting that channels showing stronger differences between vowel-evoked responses also contribute more strongly to decoding vowel memory. No significant interactions between channel selection (N-1; N-2; N-3) and vowel position and/or condition (predictable vs. random) were observed (p_FWE_ > 0.05). The second analysis did not reveal any significant effect of channel selection (main and interaction effects: p_FWE_ > 0.05; Fig. S1C), suggesting that phase coherence at the syllable rate is not related to memory decoding.

**Figure S1.**
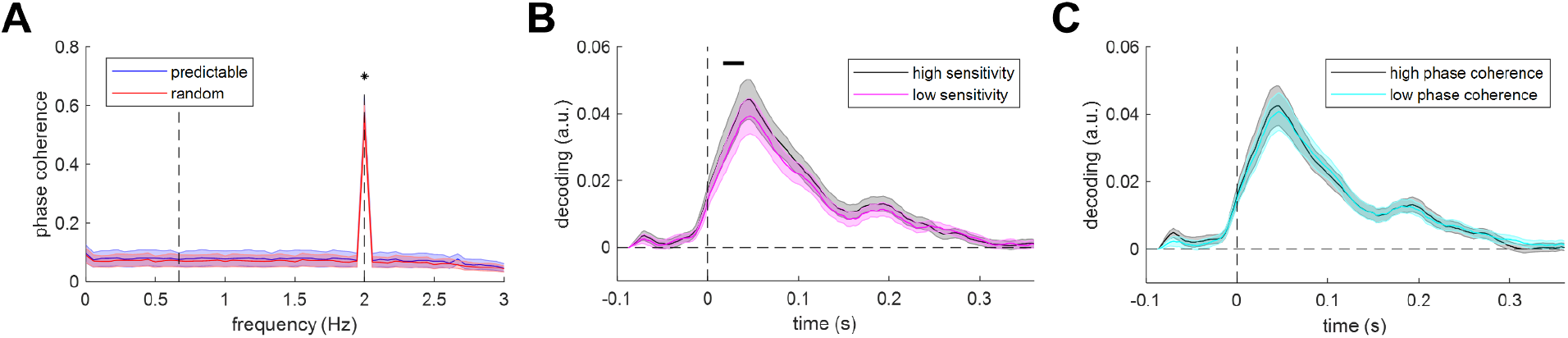
Supplementary analyses. **(A)** Phase coherence in predictable and random blocks. Asterisk denotes a significant syllable-rate peak (2 Hz). Shaded areas: SEM across recording sessions. **(B)** Decoding traces (averaged across conditions and vowel positions) based on channels showing high sensitivity to vowels (black line) vs. low sensitivity to vowels (magenta line). Horizontal bar denotes a significant difference between high vs. low sensitivity channels (p_FWE_ < 0.05). Shaded areas: SEM across recording sessions. **(C)** Decoding traces (averaged across conditions and vowel positions) based on channels showing high syllable-rate phase coherence (black line) vs. low syllable-rate phase coherence (cyan line). Shaded areas: SEM across recording sessions.

### Multivariate analysis: decoding of predicted vowels gradually improves over time

Having established that decoding of the predicted vowel (N-3) shows a specific improvement in predictable vs. random blocks, we sought to determine whether this boost shows features of a predictive representation. We reasoned that, in predictable blocks, predictions should be learned over time, and consequently the decoding of the N-3 vowel should gradually improve within and across blocks containing identical triplets (Fig. 4AE). To test this, we performed a linear regression analysis on single-trial decoding estimates, using two learning regressors - one quantifying possible gradual improvements of decoding within each sequence containing identical triplets (learning within blocks), and one quantifying possible gradual improvements of decoding over the course of the entire recording session (learning across blocks). This analysis revealed that, for the early time window in which we observed a decoding boost in the predictable vs. random condition (77-103 ms), the “within blocks” learning effect was significantly higher in the predictable than in random blocks (Wilcoxon sign rank test, Z_21_ = 2.485, p = 0.013; Fig. 4BCD), although significance testing of regression coefficients within conditions against zero did not yield significant effects (predictable: Z_21_ = 1.477, p = 0.139; random: Z_21_ = -1.825, p = 0.068). No significant learning effects across blocks were observed for the early time window (all p > 0.5). Conversely, for the later time window in which we observed a decoding boost (227-270 ms), the “across blocks” learning effect (Fig. 4FGH) was significant in the predictable condition (Z_21_ = 2.033, p = 0.042) but not in the random condition (Z_21_ = 0.122, p = 0.903), although a direct comparison of learning coefficients between conditions did not yield a significant effect (Z_21_ = 1.303, p = 0.192). The “within blocks” learning effect did not yield any significant effects in the later time window (all p > 0.5). Taken together, these results are consistent with the notion that the early N-3 decoding boost improves at faster time scales (within blocks), while the late N-3 decoding boost improves at longer time scales (across blocks).

**Figure 4.**
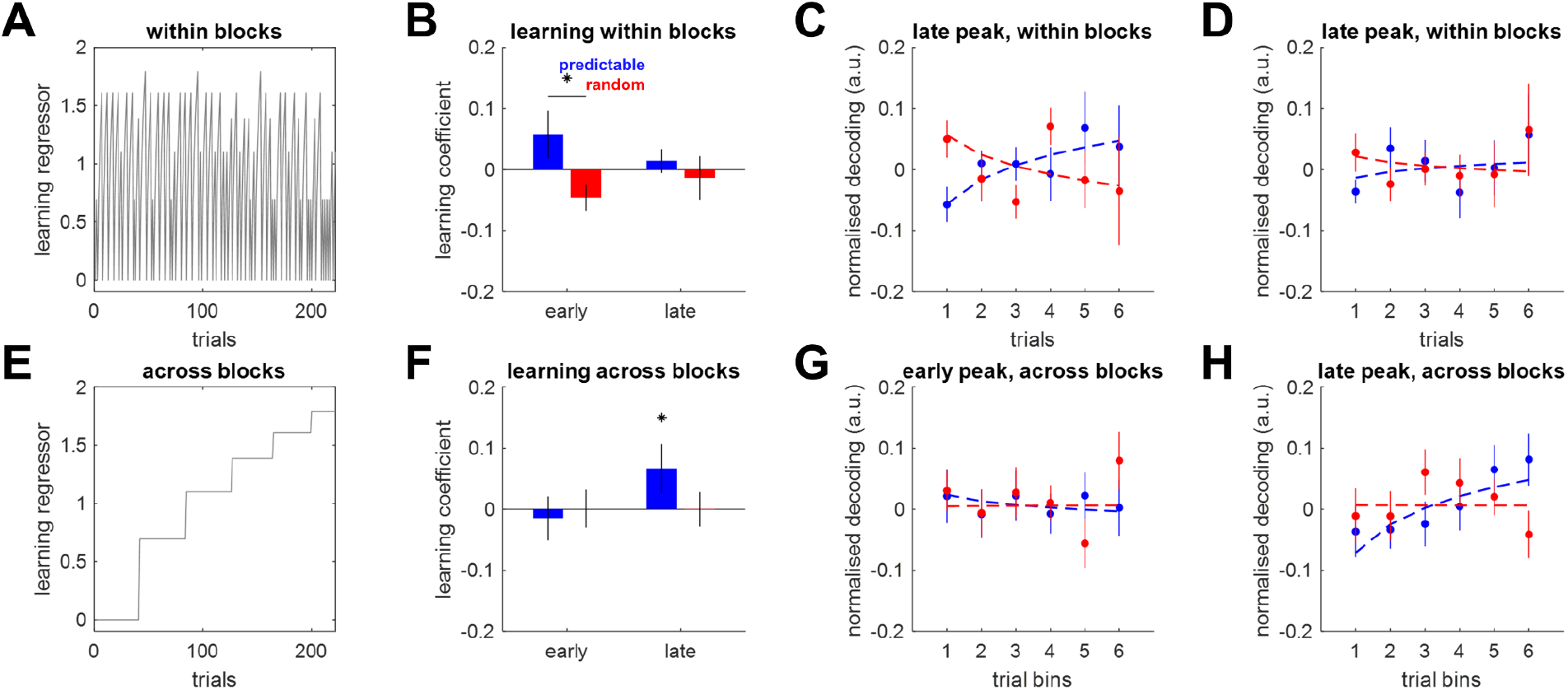
Learning effects. **(A)** A trial-by-trial regressor of learning within blocks (faster time scale) was quantified as the (log) burst number in a block of identical triplets. **(B)** Regression coefficients (“within blocks” learning) for two time windows with significant N-3 decoding boost (see Fig. 3A, right). Error bars denote SEM across recording sessions. Asterisk denotes a significant Wilcoxon sign rank test. **(C)** Normalized decoding per trial within a block of identical triplets: early time window. Error bars denote SEM across recording sessions. **(D)** Normalized decoding per trial within a block of identical triplets: late time window. **(E)** A trial-by-trial regressor of learning across blocks (slower time scale) was quantified as the block number in a recording session, binned into six bins. **(F**,**G**,**H)** Learning effects across blocks, figure legend as in (B,C,D).

### Multivariate analysis: predictive and mnemonic representations rely on independent codes

While the decoding boost observed for the N-3 vowel in predictable blocks, and its gradual improvement over time, bear the hallmarks of a predictive representation, we have also accessed mnemonic representations by decoding previous vowels (N-1 and N-2) in random blocks. To test whether these two kinds of representations rely on activity in the same neural populations in the AC, we have repeated decoding using a searchlight approach, where each decoding estimate was based on a subset of channels. We then correlated the spatial maps of decoding estimates between the predictable and random blocks. We reasoned that if predictive and mnemonic representations rely on similar populations, the decoding maps should be correlated. This analysis revealed significant correlations between spatial decoding maps in predictable and random blocks only for the N-1 vowel (Fig. 5A; 33-70 ms; t_max_ = 5.72; cluster-level p_FWE_ < 0.001), but not for the earlier vowels (N-2, N-3: all p_FWE_ > 0.05). Specifically, while for the N-1 vowel the spatial maps of decoding obtained in predictable and random blocks were almost identical and showed the strongest contribution of the anterior/inferior channels, for the N-3 vowel they were much more orthogonal, showing a inferior-superior gradient in predictable blocks and an anterior-posterior gradient in random blocks (Fig. 5BC). This pattern of results suggests that the specific decoding boost observed for N-3 vowels in predictable blocks (reflecting a predictive representation) relies on a different activity pattern than the N-3 vowel decoding in random blocks (reflecting a mnemonic representation).

**Figure 5.**
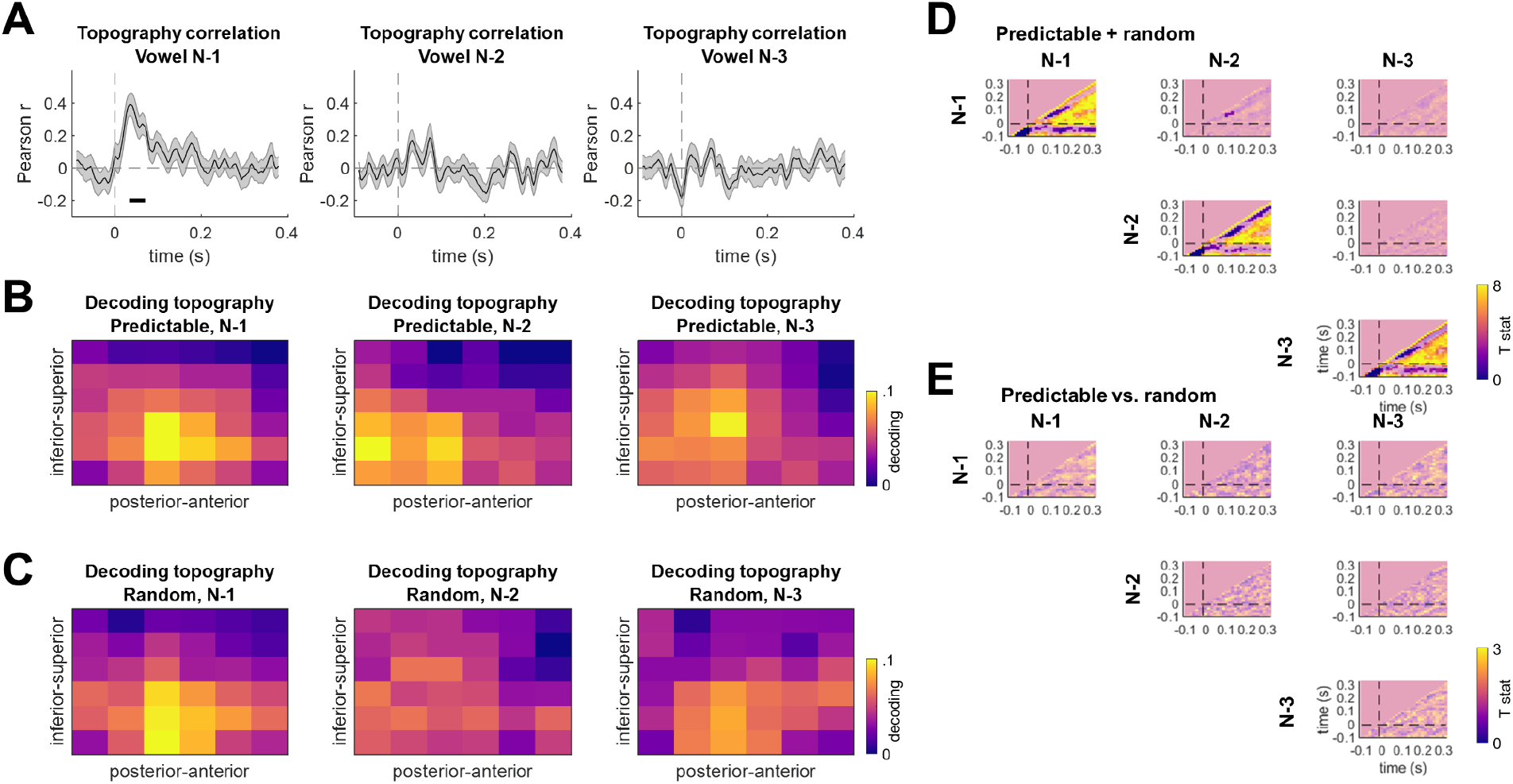
Spatial topography of predictive and mnemonic representations. **(A)** Time courses of correlation coefficients between decoding topographies in predictable vs. random blocks. Left / middle / right panel: decoding N-1 / N-2 / N-3 vowel (shaded area: SEM across recording sessions; black horizontal line: correlation coefficients significantly different from zero, p_FWE_ < 0.05). **(B)** Decoding topographies based on the 0-100 ms decoding time window, predictable blocks. Left / middle / right panel: decoding N-1 / N-2 / N-3 vowel. **(C)** Decoding topographies based on the 0-100 ms decoding time window, random blocks. Figure legend as in (B). **(D)** Cross-temporal generalization averaged across conditions (predictable vs. random). Rows: test data; columns: remaining data used for estimating decoding matrices. Each panel shows a cross-temporal decoding matrix with each time point representing decoding based on the Mahalanobis distance between a particular vowel position (N-1, N-2, N-3) and latency of neural activity and another vowel position and latency of neural activity. Unmasked areas represent significant cross-temporal decoding generalization at p_FWE_ < 0.05, cluster-level corrected. **(E)** Cross-temporal generalization: differences between conditions (predictable vs. random). Figure legend as in (D).

### Multivariate analysis: cross-temporal generalization

In a further analysis, we tested whether decoding generalizes across time points within a single response (suggesting a similar neural code for decoding based on activity measured at different latencies) and/or across vowel positions (suggesting that decoding one vowel, e.g. N-1, relies on a similar neural code as decoding another vowel, e.g. N-2). This analysis revealed that, per vowel position, decoding did generalize across time points (N-1: t_max_ = 20.21, p_FWE_ < 0.001; N-2: t_max_ = 16.05, p_FWE_ < 0.001; N-3: t_max_ = 13.64, p_FWE_ < 0.001; Fig. 5D; cf. Cappotto et al., 2021). Interestingly, we also observed negative cross-generalization between N-1 and N-2 vowels, when averaging across predictable and random conditions, such that N-2 vowel decoding at 133-167 ms post-stimulus showed impaired (negative) decoding when trained on data corresponding to the N-1 vowel at 67-100 ms (F_max_ = 25.51, t_min_ = -5.05, p_FWE_ = 0.027), possibly reflecting an interference effect between N-1 and N-2 decoding. Beyond this finding, there were no significant cross-generalization clusters between the other vowel pairs (all pFWE > 0.05), as well as no differences between predictable and random blocks either in cross-temporal generalization across time points (all pFWE > 0.05; Fig. 5E) or across vowel positions (all pFWE > 0.05). Taken together, these results suggest that decoding each triplet element relies on a relatively specific neural code, with a possible interference effect between N-1 and N-2 decoding. However, no generalization benefit could be observed for the predictable vs. random conditions.

## Discussion

In the present study, we demonstrated that stimulus history (sensory memory traces of token values up to N-3) can be decoded from neural responses to broadband noise bursts in both repeated triplet and randomized blocks, expanding on previous research employing similar decoding techniques (Cappotto et al., 2021; Wolff et al., 2020, 2015). We also demonstrate that these neural responses tap into predictive mechanisms, establishing a novel method for the decoding of both phenomena simultaneously and independent of attentional tasks. Crucially, we provide evidence for the decoding of predictive mechanisms by linking increased decodability to predictable blocks, further established through the presence of learning effects as the number of triplet pattern repeats increases. Our results also suggest that mnemonic and predictive decoding rely on largely non-overlapping neural populations.

Previous work in sensory memory decoding has established the use of broadband noise impulses in decoding recent sensory memory tokens (Wolff et al., 2020, 2015). Furthermore, our previous work in decoding auditory sensory memory tokens has established that such methods rely on mechanisms that function under anesthesia in animal models (Cappotto et al., 2021). Here, we expand on these findings by decoding further back into stimulus history, showing that it is possible to decode memory representations of both sequences and individual tokens up to N-3 using this method. We also expand on another recent study (Luo et al., 2021) showing that sequence contents can be preferentially decoded from auditory cortical activity in the rat models, but that this decoding benefit is only observed for rats that were previously trained to familiarize themselves with the sequences (rather than merely passively exposed to the sequences). Unlike the latter study, which used several interleaved sequences in a continuous stream, here we used a relatively simple stimulation protocol in which the same sequence (triplet) repeated over and over again and was then replaced with another triplet. This suggests that for sequences that are repetitive enough, decoding can be achieved even in naive and anaesthetized rats. In both the present and previous studies, we have demonstrated that univariate analysis was not sufficient to decode memory tokens and the employed multivariate methods provided significant decoding in both studies. Our findings are not likely to be explained by a simple stimulus-specific adaptation (SSA) explanation of decodability, given that we observed the decodability of randomly substituted tokens within repeated sequences as well as within non-repeating triplets. Similarly, no general adaptation effects were observed under univariate analysis, where the amplitude of tone-evoked responses did not differ between repeated and randomized trials, and burst-evoked responses were uncorrelated with vowel history, thus speaking against the idea of context differentially modulating SSA-like effects. Furthermore, if simple adaptation were responsible for decodability, this effect would be unlikely to increase with overall triplet repetition, as pattern sensitivity and resulting deviance detection has been previously shown to rely on hierarchical and contextual error detection (Casado-Román et al., 2020). Our results suggesting that decoding triplet identity and N-3 identity may occur at different latencies are consistent with the latter hypothesis.

In the present study, we demonstrate that mnemonic and predictive representations are simultaneously encoded and that neural correlates of these representations are decodable through the use of a single broadband noise burst. Predictive representations (e.g., the N-3 token in predictable blocks) are dissociable from representations of the entire triplet (see Fig. 3C) and further established via the presence of higher decodability after repeated presentations of a given triplet, implicating passive learning effects as a measure of predictive mechanisms (see Fig. 4). Interestingly, recent studies in auditory working memory observed discrete neural representations of working memory contents after the presentation of a deviant, suggesting that mnemonic template matching and working memory storage are intertwined with resultant error correction mechanisms within sensory cortices (Libby and Buschman, 2021). Although our paradigm didn’t test for this explicitly, our observation of unique topographies for mnemonic and sensory representations, and in particular the orthogonal arrangement of channel sensitivity in predictable vs. random N-3 representations, suggests that such mechanisms are not dependent on active processes and can be observed indirectly over broad neural populations even under anesthesia.

Recent studies have successfully paired concepts of statistical learning and predictive coding by investigating neural correlates of melodic expectation to naturalistic music, observing that neural responses to less statistically-likely notes elicit markers consistent with their level of statistical predictability (Di Liberto et al., 2020). Human fMRI studies in the visual domain have further established the role of temporal regularity in sequence learning and their resultant effects on the decodability of predictable stimuli (Luft et al., 2015). The simultaneous presence of mnemonic and stimulus-driven activity has also been observed in the visual domain, consistent with the predictive coding framework assumption of template matching, in that memory representations in early sensory regions can function as comparative references for incoming sensory information (Rademaker et al., 2019). Studies employing animal models and different attention states to investigate predictive coding frameworks have been lacking (Heilbron and Chait, 2018), and our results provide a clear observation of prediction formation even under anesthesia. The present literature is largely within the context of MMN responses or SSA, or employ omission designs reliant on repetition of the same stimuli token, making it difficult to separate predictive from simple adaptive mechanisms. A recent study (Parras et al., 2017) attempted to disentangle adaptive and predictive effects using an oddball paradigm in single-unit recordings from awake and anesthetized animals, demonstrating that predictive effects are organized hierarchically and suggestive of underlying MMN mechanisms. Further animal studies (Malmierca et al., 2019) have investigated neuronal pattern sensitivity with findings compatible with predictive coding frameworks reliant on temporal and spectral regularities entrained at the single-unit level, and have provided compelling evidence for a prediction-error-signaling-based explanation of hierarchical deviance detection gradient between auditory subcortical and cortical regions and prefrontal cortices (Casado-Román et al., 2020).

Our results thus fill several gaps in the animal and human electrophysiology literature. Future studies may wish to employ our approach in investigating auditory “filling in” phenomena (King, 2007) which may find an analog in our use of interrupted triplets, as such phenomena can be probed through the use of learned sequences and the resultant predictions when elements within those sequences are physically masked but perceptually and behaviorally perceived. As our findings demonstrate a correlation between statistical learning models and relative decoding strength of predictable tokens within a sequence, this technique could potentially be employed in similar naturalistic music listening paradigms to probe the effect of such models on syntactic predictability during the presentation of naturalistic stimuli such as music and language. Our searchlight approach may also allow such studies to determine if their observed effects rely on overlapping neural populations, yielding greater insights into the parallel processes of memory, learning, and prediction. The observed topographical differences in mnemonic and predictive decodability of predictable tokens may also lend important insights into the segregation of prediction-specific neuronal populations within the same hierarchically-organized cortical pathways, and future studies may wish to repeat our experiment with single-unit measurements in prefrontal and sensory cortices as well as in the hippocampus to further investigate the presence of laminar-based hierarchies implied by the predictive coding framework (Barron et al., 2020). The present study observed concurrent, yet spatially separable, mnemonic and predictable representations under anesthesia, indicating mechanisms at work in passive preparations and thus providing a new model for investigating simultaneous memory and predictive mechanisms independent of attentional state.

## Materials and Methods

### Subjects, Experimental Apparatus and Surgical Procedures

Eight young adult female Wistar rats, acquired from the Chinese University of Hong Kong, were used in the experiment. The rats were “naive”, i.e. had no experience or training with the stimulus sets prior to recording, were aged between 8 and 13 weeks (median age = 10.5 weeks), and weighed between 216 and 289 g (median weight = 238 g). Normal hearing was ascertained by measuring auditory brainstem response at thresholds < 20 dB sound pressure level (SPL) to broadband click trains. Anaesthesia was induced with an intraperitoneal (i.p.) injection of ketamine (80 mg/kg) and xylazine (12 mg/kg), and maintained throughout the experiment via 20% urethane injections. A first dose of 0.25 ml/kg of the urethane solution was administered one hour after the induction with ketamine and xylazine, and further 0.25 ml/kg doses were delivered as required, based on periodic assessments of anesthesia depth via the toe pich withdrawal reflex. Dexamethasone (0.2 mg/kg, i.p.) was delivered before surgery as an anti-inflammatory. This protocol, based on previous rodent studies (Cappotto et al., 2021; Malmierca et al., 2019), allowed for fast induction of anesthesia via the initial administration of ketamine and xylazine, while avoiding later NMDA-specific inhibitory effects of ketamine through the use of urethane to maintain anesthesia for ECoG recordings. The anaesthetized animal was placed in a stereotaxic frame, and the animal’s head was fixed with hollow ear bars to allow sound delivery. An isothermal heating pad and a rectal thermometer were used to maintain body temperature at 36 ± 1°C throughout the experiment. The skin and muscle tissue over the temporal lobe of the skull were removed, and a craniotomy was performed to expose a 5×4 mm region over the right AC, leaving the dura intact. The anterior edge of the craniotomy was 2.5 mm posterior from Bregma, and the dorsal edge was 2 mm ventral from Bregma (Fig. 1A, adapted from Cappotto, et al., 2021). The ECoG array was placed on the exposed cortex and a cotton roll was placed between the remaining skin and the array to hold the array securely in place and ensure a stable, low impedance contact between the recording sites and the dura. A hole was drilled through the skull anterior to the Bregma on the animal’s left to place a small stainless steel screw which served as ground and reference electrode for the electrode array and headstage amplifier.

Correct placement of the ECoG array was verified by recording a set of Frequency Response Areas (FRAs) from each site by collecting responses to 100 ms pure tones varying in sound level (30 - 80 dB SPL) and frequency (500 - 32,000 Hz, ¼ octave steps). Each tone was presented 10 times, in a randomly interleaved fashion, with an onset-to-onset ISI of 500 ms.

### Experimental Paradigm and Stimulus Design

The artificial vowels were generated using custom Python scripts. These generated pulse trains which were subsequently passed through a cascade of two 2nd-order Butterworth bandpass filters with a bandwidth equal to 20% of the center (formant) frequency (scipy.signal functions butter() and lfilter()). The formant frequencies for these artificial vowels were chosen to lie between 900 and 9000 Hz to bring them well into the auditory range of rats, and the fundamental frequencies (F0s) of the vowels were relatively low, between 260 and 420 Hz, to generate a large number of closely stacked harmonics under each formant. Stimulus sequences consisted of combinations of three possible artificial vowels, one we refer to as “A” with formants and 3000 and 5400 Hz and an F0 of 420 Hz, an “O” with formants 900 and 2700 Hz and F0 260 Hz, and an “I” with formants 1050 and 9000 Hz and F0 300 Hz. On occasion, as described further below, one of the vowels in the sequence could be replaced by either a 150 ms frozen pink noise burst computed according to the algorithm described in https://github.com/python-acoustics/python-acoustics/blob/master/acoustics/generator.py, or by a silent pause. The artificial vowel and pink noise tokens were loaded onto a Tucker Davis Technologies (TDT) RZ6 digital sound processor which was programmed using custom written software to present the tokens in a predefined order at a sample rate of 48,828 Hz through headphone drivers connected to the hollow ear bars via 3D printed adapters.

Two types of blocks were employed. In “predictable” blocks, vowels were grouped into one of six triplets (AAO, AOO, AAI, AII, OOI, OII) with each triplet presented at least 25 times in a given block of identical triplets before being replaced with another triplet. Each session contained ∼72 such blocks (amounting to a total of 2100 triplets per session), presented in a different order per session. Triplets were selected to prevent redundant combinations from occurring during presentation (e.g., AOO, OAO, and OOA would result in identical sequences with different starting points, and thus only AOO was used). The triplets were then concatenated to form the long stimulus sequences presented in the experimental sessions. In these sequences, 5% of stimulus events were replaced with omissions, and 5% were similarly replaced with a burst of pink noise. The vowels that were replaced with noise bursts or silent pauses were chosen pseudo-randomly, subject to the constraint that a minimum of three repetitions of a given triplet had to have occurred before a vowel could be replaced. In a control condition (“random” sessions), vowels were presented randomly, rather than in predefined triplets. The positions of omissions and noise bursts within the stimulus sequences were kept the same across the predictable and random blocks.

### Neural data acquisition and pre-processing

An 8 × 8 Viventi ECoG electrode array with 400 µm electrode spacing (Woods et al., 2018) was used to acquire ECoG recordings, employing three ground channels located in the corners of the array, and a common reference. A (TDT) PZ5 neurodigitizer was used to record signals from the array via a RZ2 processor. FRA responses were recorded with BrainWare software at a sampling rate of 24,414 Hz, and responses to the vowel sequences were recorded using custom Python code at a sampling rate of 6104 Hz. The recorded electrode signals were first low-pass filtered at a cutoff frequency of 90 Hz using a 5th order Butterworth filter, and downsampled to 300 Hz to extract neural activity evoked by acoustic stimuli. The pre-processed signals were re-referenced to the average of all channels (Ball et al., 2009), and segmented by extracting 500 ms long voltage traces from -100 ms to +400 ms relative to the onset of each token. Epoched traces were baseline-corrected by subtraction of the mean pre-stimulus voltage values, and linearly detrended (Salisbury, 2012).

### Univariate analysis: summarising vowel-evoked, omission-evoked, and frozen noise burst-evoked activity

Univariate analysis was performed to assess whether vowel types (A, I, O) modulated vowel-evoked, burst-evoked, and omission-evoked activity on a channel-by-channel basis. Additionally, in the analysis of burst-evoked and omission-evoked activity, we tested whether it is modulated by the preceding sounds at different “positions” relative to the burst/omission (N-1 position: the immediately preceding vowel, N-2 position: two stimuli before the burst/omission, N-3 position: three stimuli before the burst/omission). Epoched data were separated per vowel, position, and condition, and then averaged across trials. First, to visualise the evoked responses, trial-averaged ECoG responses were concatenated across sound types/positions/conditions/animals, resulting in 2 two-dimensional matrices per condition with single channels along one dimension and concatenated time points along the second dimension. A principal component analysis using singular value decomposition was performed on the resulting matrices. The output provided spatial principal components describing channel topographies, and temporal principal components describing voltage time-series concatenated across vowels/positions and animals, sorted by the ratio of explained variance. A weighted average was calculated to summarize the top principal components explaining 95% of the original variance, weighted by the proportion of variance explained. These resulting voltage time-series were averaged per vowel across animals. Frozen noise burst-evoked and omission-evoked single-trial data were similarly averaged across trials, separately for each preceding vowel and position, and subject to the same principal component analysis described above.

The above principal component analysis was used only for the purposes of visualizing the data. In order to test if any time points and channels showed significant amplitude modulations by vowel (in case of vowel-evoked responses) or preceding vowel in each position (in case of burst-evoked and omission-evoked responses), single-subject trial-average ECoG data in the original electrode grid were converted into three-dimensional matrices containing two spatial dimensions and one temporal dimension. These matrices were then converted to 3D images and entered into a repeated-measures ANOVA with one within-subjects factor (vowel; three levels) and one repeated-measures factor (rat), implemented in SPM as a general linear model (GLM). This was done separately for each stimulus type (vowel-evoked responses, burst-evoked responses, and omission-evoked responses). The effects of preceding vowels on burst-evoked and omission-evoked responses were analyzed in separate ANOVAs per position. To test for the effect of vowel on evoked activity amplitude, an omnibus F test across 3 vowels was used. The resulting statistical parametric maps were thresholded at p < 0.005 (two-tailed) and corrected for multiple comparisons across spatiotemporal voxels at a family-wise error (FWE)-corrected p_FWE_ = 0.05 (cluster-level) (Kilner et al., 2005).

### Univariate analysis: oscillatory activity

To test whether sequence processing is associated with spectral peaks in the neural response spectrum at the syllable and triplet rate (Henin et al., 2021), we analyzed phase coherence of neural activity. Specifically, for each rat and recording session, we split the continuous single-channel ECoG data into 175 chunks of 12 triplets, and, for each chunk, calculated the Fourier spectrum of neural activity measured during that chunk. Inter-trial phase coherence (ITPC) was calculated according to the following equation (Ding and Simon, 2013):

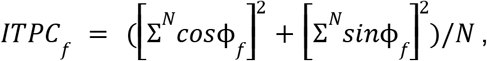

where *φ*_*f*_ denotes the Fourier phase at a given frequency *f* and *N* = 175 chunks. In the initial univariate analysis, phase coherence estimates were averaged across channels. To test for the presence of statistically significant phase coherence peaks, coherence values at the token rate (2 Hz) and triplet rate (0.667 Hz) were compared against the mean of coherence values at their respective neighbouring frequencies (single token rate: 1.944 and 2.056 Hz; triplet rate: 0.611 and 0.722 Hz) using Wilcoxon’s signed rank tests.

### Multivariate analysis: decoding sensory, mnemonic, and predicted vowel information

Data were subjected to multivariate analyses to test if information about vowel type could be decoded from the pattern of burst-evoked and omission-evoked activity observed across multiple channels and time points. To this end, we adapted methods established in previous multivariate decoding research, which has demonstrated decodability in similar data and experimental contexts (Cappotto et al., 2021; Myers et al., 2015; van Ede et al., 2018; Wolff et al., 2020, 2017).

Prior to decoding, single-trial omission or frozen noise burst-evoked responses were sorted by the preceding vowel, separately for each vowel position. Decoding time-courses were estimated using a sliding window approach (Cappotto et al., 2021; Wolff et al., 2020), pooling information over multiple time-points and channels to boost decoding accuracy (Grootswagers et al., 2017; Nemrodov et al., 2018). Specifically, for each channel, trial, and time point, we first pooled voltage values within a 50 ms window relative to a given time point. Then, a vector of 5 average voltage values was calculated per channel and trial by downsampling the voltage values over 10 ms bins. The data were then de-meaned to remove the channel-specific average voltage over the entire 50 ms time window from each channel and time bin, ensuring that the multivariate analysis approach was optimized for decoding transient activation patterns (Cappotto et al., 2021; Wolff et al., 2020). For the subsequent leave-one-out cross-validation decoding, the vectors of binned single-trial temporal data were then concatenated across channels. We used the Mahalanobis distance (De Maesschalck et al., 2000) as a multivariate decoding metric to take advantage of the potentially monotonic relation between vowel category and neural activity (Auksztulewicz et al., 2019; Cappotto et al., 2021; Wolff et al., 2020). Responses to dissimilar vowels are expected to yield large Mahalanobis distance metrics, while responses to similar vowels are expected to yield low Mahalanobis distance metrics. Having been shown to be optimal for decoding (Grootswagers et al., 2017), a leave-one-out cross-validation approach was used per trial, wherein we calculated 3 pairwise distances between ECoG amplitude fluctuations measured in a given test trial and mean vectors of ECoG amplitude fluctuations averaged for each of the 3 vowels/positions in the remaining trials. A shrinkage-estimator covariance obtained from all trials, excluding the test trial, was used to compute the Mahalanobis distances (Ledoit and Wolf, 2004). Combining Mahalanobis distance with Ledoit–Wolf shrinkage has been shown to have performance advantages over other correlation-based methods of measuring brain-state dissimilarity (Bobadilla-Suarez et al., 2019), while Mahalanobis distance-based decoding has known advantages over linear classifiers and simple correlation-based metrics (Walther et al., 2016).

Single-trial relative Mahalanobis distance estimates were averaged across trials, resulting in a 3 × 3 distance matrix for each rat, time point, relative vowel position (N-1, N-2, N-3), and substitution type (noise vs. omission). To obtain overall decoding quality traces, the 3 × 3 distance matrices were subject to a subtraction of the averaged off-diagonal elements (mean distance between vowels) from the averaged diagonal elements (mean distance within vowels). The resulting decoding time-series were entered into a 2×3 repeated-measures ANOVA with within-subjects factors Block (predictable vs. random) and Position (N-1, N-2, N-3), separately for the two substitution types (noise vs. omission). The resulting statistical parametric maps were thresholded at p < 0.005 (uncorrected). Across time points, p values were corrected using a family-wise error approach at a cluster-level p_FWE_ = 0.05 (Kilner et al., 2005).

We reasoned that significant decoding of the N-3 vowel in the predictable blocks, but not in the random blocks, would reveal predictive representations of the expected vowel. In a follow-up analysis, we wanted to test bursts/omissions reactivate representations containing (1) information about the entire preceding triplet, or (2) specific information about the N-3 vowel, independent of the rest of the triplet. To this end, we ran an additional decoding analysis, this time using a 18 × 18 stimulus matrix (corresponding to 18 possible triplets, with 3 phase shifts for each of the 6 unique triplets; e.g., for a unique triplet AAO, the three phase shifts would correspond to AAO, AOA, and OAA), yielding 18 × 18 Mahalanobis distance matrices. This analysis focused on the predictable blocks only and zoomed into two time clusters in which we observed significant N-3 vowel decoding (see Results). To quantify the decoding of the entire triplet, we subtracted the mean of all off-diagonal elements of the 18 × 18 stimulus matrix from the mean of all diagonal elements (Fig. 3C). To quantify the decoding of information about the N-3 vowel independent of the entire triplet identity, we subtracted the mean of those elements of the 18 × 18 stimulus matrix which did not share the first vowel from the mean of those elements of the matrix which did share the first vowel (excluding the diagonal elements, corresponding to identical triplets). The resulting decoding estimates were subject to one-sample t-tests (two-tailed) across recording sessions.

In an additional analysis, since we observed univariate differences in vowel-evoked responses (see Results), we tested whether decoding primarily relies on those channels that are also associated with sensory encoding of vowels. To this end, we repeated the decoding analysis for two subsets of channels - those which strongly differentiated between vowels (with the corresponding F statistic of the main effect of vowel on the vowel-evoked responses higher than the median across channels) and those which differentiated weakly between vowels (F statistic below median across channels). The resulting decoding time-series were compared between the two groups of channels using a series of paired t-tests, correcting for multiple comparisons across time points at a family-wise error (FWE)-corrected p_FWE_ = 0.05 (cluster-level) (Kilner et al., 2005).

While we did not observe univariate differences in spectral peaks at the single vowel rate between condition (and we did not observe peaks at the triplet level overall; see Results), in a further analysis we also tested whether decoding might rely on those channels which show relatively higher triplet-rate peaks than other channels. Again, we repeated the decoding analysis for two subsets of channels, this time splitting them based on the single-channel phase coherence estimates for the single vowel rate (2 Hz; above/below median). The two resulting decoding time-series were compared using a series of paired t-tests, correcting for multiple comparisons as above.

### Multivariate analysis: learning effect on decoding

Another question we wanted to address is whether any decoding benefit we might observe in the predictable stimulus condition reflects predictive neural processing. In particular, we hypothesized that, if the decoding boost in predictable blocks is related to predictive processing, it should gradually build up, as the auditory system needs time to detect repeating patterns and learn to use them for predictions of which sound token is expected when. This can occur at two time scales: first, decoding can improve with each subsequent vowel token embedded in a block of identical triplets (reflecting learning within blocks); second, decoding can improve over subsequent blocks (reflecting learning across blocks). To test these hypotheses, we constructed two trial-by-trial learning regressors - a “within blocks” regressor quantifying the vowel position within a block of identical triplets, and an “across blocks” regressor quantifying which block of a particular triplet it is within the entire recording session. To facilitate comparisons between the two regressors, the “within blocks” regressor only included vowel position from 1 (first burst within a sequence) to 6 (sixth burst), while the “across blocks’’ regressor was binned into 6 bins of 2 blocks in each bin (e.g., bin 1 contained the first 2 blocks of a particular triplet, while bin 6 contained the last 2 blocks of the same triplet). Both regressors were log-transformed to increase the relative effect of the first bursts/sessions relative to the last bursts/sessions (HiJee et al., 2021). We then repeated the decoding analysis of the N-3 vowel and, per recording session and condition, performed a multiple linear regression with a constant term and the two learning regressors on single-trial decoding estimates. Specifically, for both of the time clusters in which we identified significant differences between decoding in predictable vs. random blocks, we selected the single-trial peak decoding within a given time cluster, and then normalized (z-scored) the trial-by-trial peaks per rat, recording session, and condition. This resulted in 8 sets of learning coefficients: (1) for predictable vs. random conditions, (2) quantifying learning within vs. across blocks, (3) estimated for early vs. late time window. The resulting regression coefficients (betas) were tested for significant differences between predictable and random blocks, as well as for significant differences against zero (separately for each block), using Wilcoxon sign rank test.

### Multivariate analysis: similarity between predictive and mnemonic representations

To test whether the predictive and mnemonic representations are shared, we quantified the spatial correlation of decoding topographies between predictable and random blocks. Our reasoning was that, if predictive and mnemonic representations are shared, decoding topographies should be similar between predictable and random blocks. On the other hand, if predictive and mnemonic representations are independent, the decoding topographies should be different between the two types of blocks. To this end, we repeated the decoding analysis, this time using a searchlight approach. Specifically, rather than using all channels for decoding, we used subsets of channels, with each subset forming a 3×3 grid. Different subsets overlapped by 1 row or column, resulting in 36 (6×6) decoding estimates based on the 3×3 grids, separately for each recording session, condition, and time point. We then correlated the spatial maps obtained for predictable and random blocks, separately for each recording session and time point. The resulting Pearson correlation coefficients were entered into a series of one-sample t-tests, correcting for multiple comparisons across time points at p_FWE_ = 0.05 (Kilner et al., 2005).

### Multivariate analysis: cross-temporal generalization

In a further analysis, we tested whether decoding a particular vowel generalizes across time points (suggesting that the reinstated representations rely on a similar neural code, independent of the latency of measured neural activity) and/or across vowel positions (suggesting that decoding one triplet element relies on a similar neural code as decoding another triplet element). To this end, we performed a cross-temporal generalization analysis, in which we repeated our multivariate decoding analysis but with an important modification of the leave-one-out cross-validation approach. First, to quantify generalization across time points, in calculating the Mahalanobis distance we incrementally shifted the latency of the test data with respect to the remaining trials, in 16 ms time steps - such that decoding was trained on one latency but tested on another. As a result of this approach, rather than decoding time series, per recording session and condition (predictable vs. random) we obtained decoding matrices with each matrix element representing the Mahalanobis distance between data measured at two different latencies. Second, to quantify generalization across vowel positions, we allowed the test data labels to be replaced by labels corresponding to another vowel than the remaining trials. As a result of this approach, rather than obtaining 3 decoding matrices (one per vowel position), we obtained 6 decoding matrices with the 3 additional matrices representing the Mahalanobis distance between data measured at two different vowel positions. The resulting decoding matrices were entered into a series of 6 GLMs (one per vowel position pair), each implementing a paired t-test between decoding estimates obtained for the predictable and random conditions. The resulting statistical parametric maps were thresholded at p < 0.005 (two-tailed) and corrected for multiple comparisons across spatiotemporal voxels at a family-wise error (FWE)-corrected p_FWE_ = 0.05 (cluster-level) (Kilner et al., 2005).

## Acknowledgments

This work has been supported by the European Commission’s Marie Skłodowska-Curie Global Fellowship (750459 to R.A.), the Hong Kong General Research Fund (11100518 to R.A. and J.S.) and a grant from the European Commission/Hong Kong Research Grants Council Joint Research Scheme (9051402 to R.A., D.P. and J.S.).

